## Supplementary for "Surface marker expression in small and medium/large mesenchymal stromal cell-derived extracellular vesicles in naïve or apoptotic condition using orthogonal techniques"

### Supplementary Methods

Apoptosis assay

Cytofluorimetric evaluation of apoptotic cells was performed using the Muse™ Annexin V & Dead Cell Kit (Merck-Millipore, Burlington, MA, USA), according to the manufacturer's instructions. Briefly, $10 \times 10^3$ naïve cells or cells were starved with RPMI medium with or without addition of 500ng of anti-Fas targeted antibody for 6, 16 or 24h. Cells were then detached and resuspended in Muse™ Annexin V & Dead Cell Kit (Luminex, Austin, TX, USA), and the percentage of apoptotic cells (Annexin V+) was detected.

Flow cytometry

Flow cytometry analysis of the ApoBDs was performed using BD FACSCelesta™ Flow Cytometer. ApoBDs were compared with apoptotic MSCs cells and latex beads of size 4 µm stained with FITC Annexin V antibody for 30min at room temperature.

### Supplementary Figures

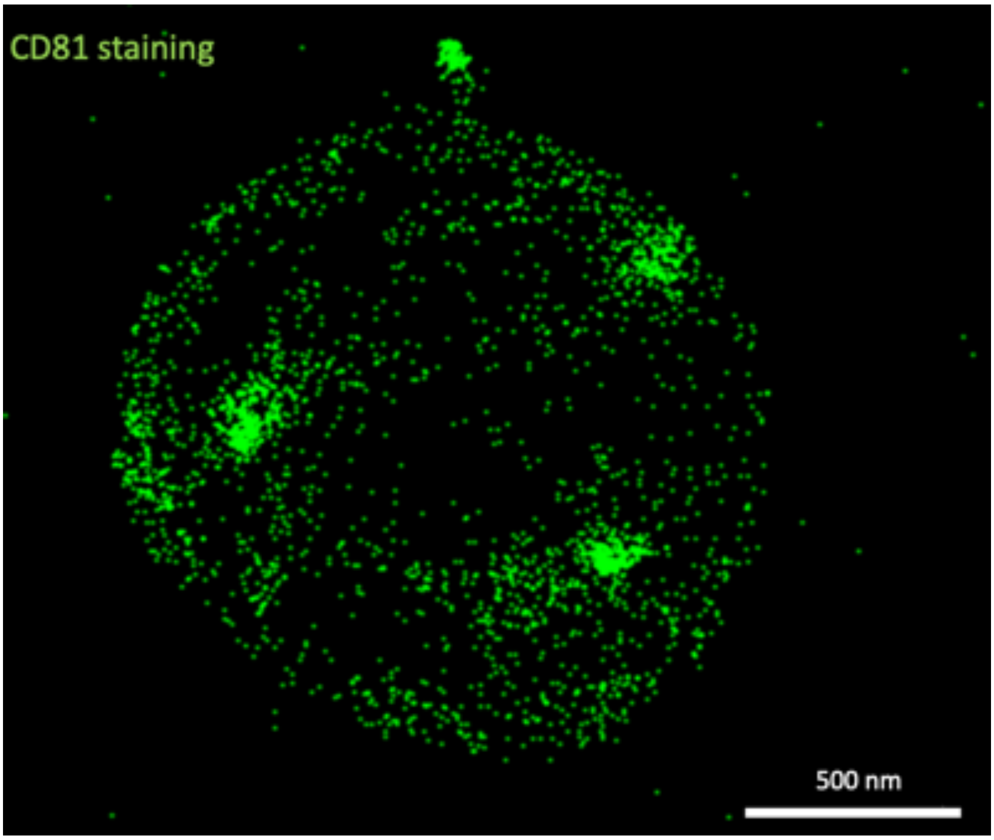


Supplementary figure 1: Representative super-resolution microscopy images of a single EV from the 10k BM-MSC fraction, showing CD81 tetraspanin surface distribution along the membrane, with areas of condensed expression.

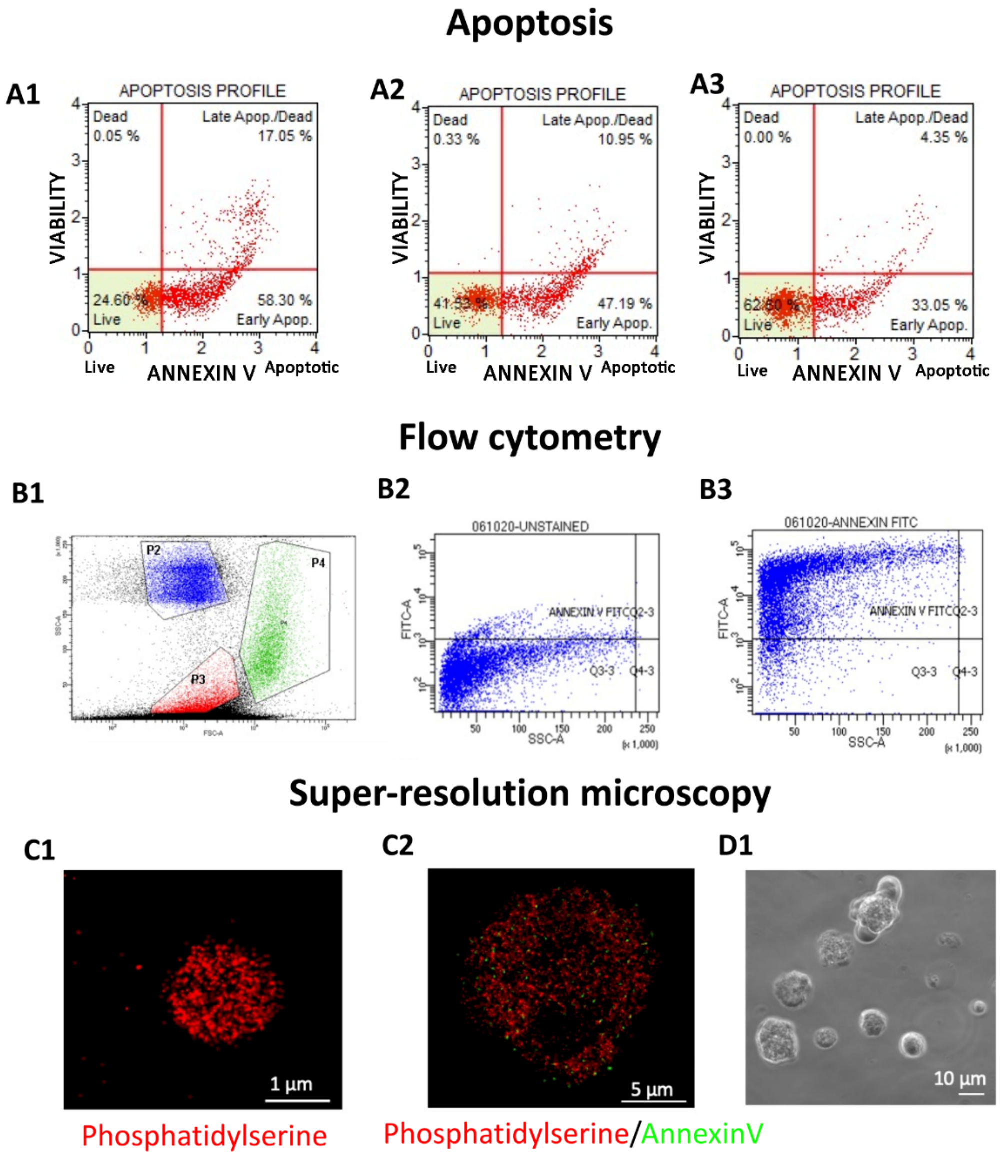


Supplementary figure 2: Induction of apoptosis and characterization of apoptotic bodies using MUSE flow cytometry and super-resolution microscopy. (A) MSCs viability profile after 24h in RPMI with 500ng of anti-Fas antibody (A1), RPMI (A2) and AlphaMEM with 10%FBS (A3). (B) Representative images of flow cytometric analysis showing Annexin V expression. (B1) flow cytometry gating strategies, P4: apoptotic bodies (ApoBDs), P2: 4 µm latex beads, P3: apoptotic cells. Representative images of flow cytometric analysis showing (B2) unstained ApoBDs and (B3) ApoBDs stained with Annexin V. (C) Representative super-resolution microscopy image of ApoBD fraction stained (C1) with Annexin V (red), (C2) with Annexin V (red) and Phosphatidylserine (green). (D1) Representative bright field microscopy image of ApoBDs realised by MSCs undergoing apoptosis.

### Supplementary Table

Supplementary Table 1: MUSE assay used to set up the apoptotic induction. Cells were analysed after 6, 16 and 24h in full condition medium, RPMI or RPMI with 500ng of anti-Fas antibody. Graphical visualization is in supplementary Fig. 2A.

| 6h | AlphaMEM + 10% FBS | RPMI | RPMI+ anti-Fas | SD |
|---|---|---|---|---|
| Live | 80.45% | 71.94% | 72.03% | 0.07 |
| Early apoptosis | 15.95% | 23.38% | 24.75% | 0.07 |
| Late apoptosis | 3.50% | 4.28% | 3.03% | 0.01 |

| 16h | AlphaMEM 10% FBS | RPMI | RPMI+ anti-Fas | SD |
|---|---|---|---|---|
| Live | 88.69% | 81.99% | 59.92% | 0.01 |
| Early apoptosis | 7.07% | 13.25% | 17.58% | 0.02 |
| Late apoptosis | 3.89% | 4.55% | 20.95% | 0.00 |

| 24h | AlphaMEM 10% FBS | RPMI | RPMI+ anti-Fas | SD |
|---|---|---|---|---|
| Live | 62.60% | 41.53% | 24.15% | 0.01 |
| Early apoptosis | 33.05% | 47.19% | 57.70% | 0.01 |
| Late apoptosis | 4.35% | 10.95% | 18.13% | 0.02 |
